# Comparative analysis reveals the modular functional build-up of megaplasmid pTTS12 of *Pseudomonas putida* S12: a paradigm for transferable traits, plasmid stability and inheritance?

**DOI:** 10.1101/2020.06.19.162511

**Authors:** Hadiastri Kusumawardhani, Rohola Hosseini, Johannes H. de Winde

**Affiliations:** Institute of Biology Leiden, Leiden University, The Netherlands

**Keywords:** *Pseudomonas putida*, genome sequence, solvent tolerance, megaplasmid, mobile genetic elements

## Abstract

The *Pseudomonas putida* S12 genome contains 583 kbp megaplasmid pTTS12 that carries over 600 genes enabling tolerance to various stress conditions, including the solvent extrusion pump SrpABC. We performed a comparative analysis of pTTS12 against 28915 plasmids from NCBI databases. We investigated putative roles of genes encoded on pTTS12 and further elaborated on its role in the establishment and maintenance of several stress conditions, specifically focusing on solvent tolerance in *P. putida* strains. The backbone of pTTS12 was found to be closely related to that of the carbapenem-resistance plasmid pOZ176, member of the IncP-2 incompatibility group, although remarkably the carbapenem resistance cassette is absent from pTTS12. Megaplasmid pTTS12 contains multiple transposon-flanked cassettes mediating resistance to various heavy metals such as tellurite, chromate (Tn7), and mercury (Tn5053 and Tn5563). Additionally, pTTS12 also contains a P-type, Type IV secretion system (T4SS) supporting self-transfer to other *P. putida* strains. This study increases our understanding in the build-up of IncP-2 plasmids and several promising exchangeable gene clusters to construct robust microbial hosts for biotechnology applications.

**Importance:** Originating from various environmental niches, large numbers of bacterial plasmids have been found carrying heavy metal and antibiotic resistance genes, degradation pathways and specific transporters for organic solvents or aromatic compounds. Such genes may constitute promising candidates for novel synthetic biology applications. Our systematic analysis of gene clusters encoded on megaplasmid pTTS12 underscores that a large portion of its genes is involved in stress response increasing survival under harsh conditions like heavy metal and organic solvent resistance. We show that pTTS12 belongs to the IncP-2 plasmid family. Comparative analysis of pTTS12 provides thorough insight into the structural and functional build-up of members of the IncP-2 plasmid family. pTTS12 is highly stable and carries a complex arrangement of transposable elements containing heavy metal resistance clusters as well as distinct aromatic degradation pathways and solvent-extrusion pump. This offers interesting insight into the evolution of solvent tolerance in the *P. putida* family.

## Introduction

### Megaplasmids as transferable vehicles of environmental resistance genes

Bacteria may use plasmids as autonomous, self-replicating elements driving horizontal transfer of genes (HGT) that confer resistance to otherwise detrimental conditions. As such, these extrachromosomal entities often confer advantageous characteristics for the host strain (1–3). The rapid spread of resistance genes through these, usually large conjugative plasmids is further facilitated by the presence of a variety of mobile genetic elements such as transposons, integrons and insertion sequences (IS’s) (4–7). The spread of multidrug resistance (MDR) via mobile genetic elements has been the subject of investigation for a number of years (8). Despite those efforts, the role and functioning of megaplasmids is still poorly understood. Recent studies highlighted the role of megaplasmids in the spread of MDR in the opportunistic pathogen *Pseudomonas aeruginosa* (1, 2). Strains of its close relative, *Pseudomonas putida*, have been known to harbor large conjugational plasmids, conferring resistance to environmental threats (3, 9). Plasmids of the *Pseudomonas* family are classified by incompatibility groups, that exhibit various modes of compatibility and transferability (10). We have chosen to study the recently identified megaplasmid pTTS12 of *Pseudomonas putida* S12 in comparison with a number of other large plasmids of the *Pseudomonas* family, in order to elucidate key elements governing HGT, stability and essential functions.

### Solvent tolerance in *Pseudomonas putida* S12; resistance as a benefit for biotechnology

*Pseudomonas putida* S12 is a gram-negative soil bacterium which was isolated on styrene as a sole carbon source (11). This strain shows a remarkable tolerance towards non-metabolized organic solvents (e.g. toluene) (12). Such high tolerance towards organic solvents presents a beneficial trait and advantage for bioproduction of aromatics and biofuel (13, 14). Due to its solvent tolerance and versatile metabolism, *P. putida* S12 excels as a microbial host for production of valuable chemicals (15–19). Removal of organic solvent molecules from the bacterial cell membrane is essential and carried out by SrpABC, a resistance-nodulation-cell division (RND) family efflux pump (12, 20). Membrane compaction and the upregulation of chaperones, general stress responses, and TCA cycle-related enzymes support a further intrinsic solvent tolerance in *P. putida* S12 (21–24).

We recently found through whole-genome sequencing that the genome of *P. putida* S12 consists of a 5.8 Mbp chromosome and a 583 kbp single-copy megaplasmid pTTS12 (25). Interestingly, both the SrpABC RND-efflux pump and the styrene degradation pathway which are major distinctive features of *P. putida* S12, are encoded on this megaplasmid. The genome of *P. putida* S12 contains a large number of several types of mobile elements, spread over both chromosome and megaplasmid pTTS12 (25–27). Some of these insertion sequences were shown to be involved in the regulation and adaptation towards stress conditions, for example during the solvent stress (4, 28). Like *P. putida* S12, related solvent-tolerant *P. putida* DOT-T1E and *Pseudomonas taiwanensis* VLB120 have been shown to harbor megaplasmids of 121 kb and 312 kb, respectively (29, 30). Indeed, those plasmids encode RND-type efflux pumps and biodegradative pathways for aromatic compounds, similar to *P. putida* S12. To further characterize pTTS12, we here performed a comparative analysis against a large number of other megaplasmids from Refseq and Nuccore databases. With this analysis, we aimed at identifying the origin of the pTTS12 plasmid and understanding the build-up of environmental-stress related gene clusters in pTTS12.

## Results

### Comparative analysis of megaplasmid pTTS12

The 583 Kbps megaplasmid pTTS12 (GenBank accession no. CP009975) is a single-copy plasmid encoded in *P. putida* S12 (25). For this paper, annotations for pTTS12 were further extended (Table S1) and the overall sequence was corrected based on our previous observations of an additional seven ISS12 mobile elements (4). pTTS12 encodes 609 genes of which 583 are single copy and 15 genes are duplicated at least once. These genes were divided into 232 putative operons for subsequent ordered genes in forward or reverse strand with less than 20 bps distance. The operons were clustered together based on the function of genes encoded within these clusters (Table S1).

Comparative analysis was performed with 28915 plasmids larger than 2 kb in length, acquired from the Refseq database (https://www.ncbi.nlm.nih.gov/refseq/) combined with additional sequences from the Nuccore database (https://www.ncbi.nlm.nih.gov/nuccore/). The top 50 plasmids encoding homologues of pTTS12 proteins are visualized in a circular plot using CGView (31) as shown on Figure 1, and listed in Table 1. In Figure 1, the outer purple line represents pTTS12 with forward coding genes on top of the line and the reverse coding gene sequences below the line. Each of the subsequent 50 inner circles represents a single plasmid and is colored based on the family of its host. In case no homologous protein was identified for a pTTS12 counterpart, that space on the circle was left blank. The extended circular plot and list of the top 500 plasmids encoding homologues of pTTS12 can be found in Figure S1 and Table S2.

**Table 1.**
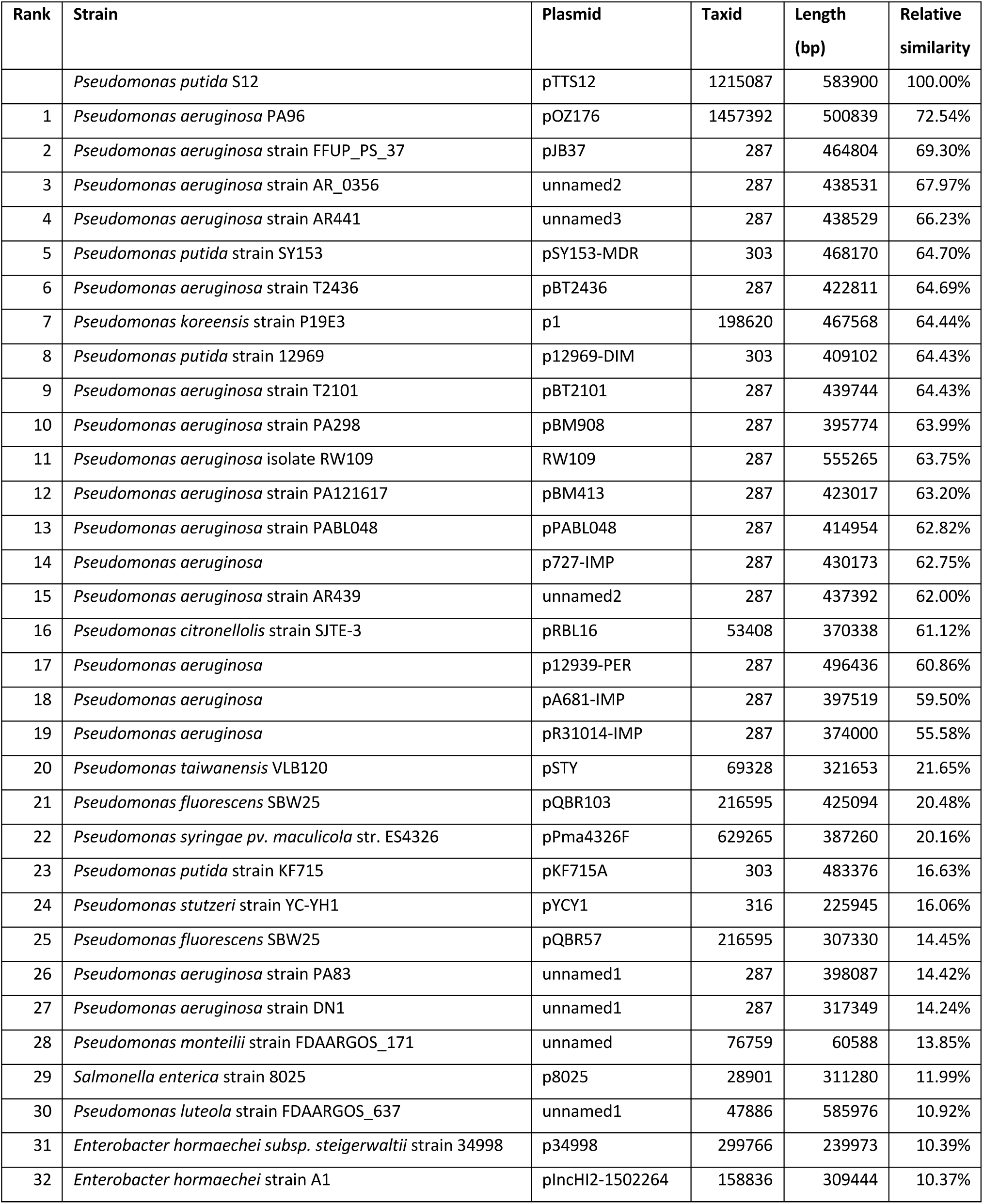
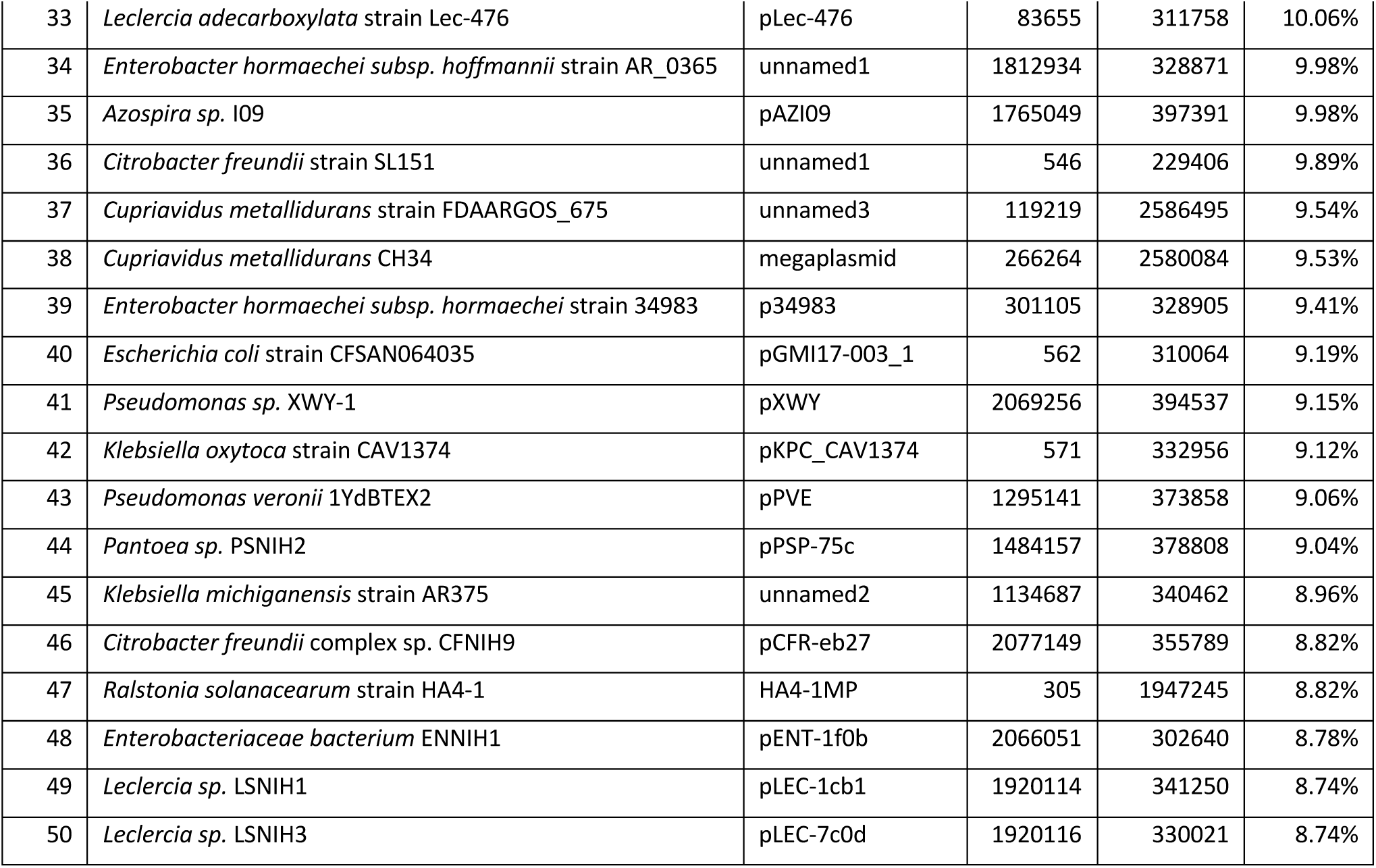
List of 50 plasmids with the highest similarity scores to pTTS12. pTTS12 coding sequences (CDS) were aligned to other megaplasmids available at NCBI databases. The relative similarity score was calculated by dividing total scores for each plasmid by total score obtained for pTTS12 itself.

**Figure 1.**
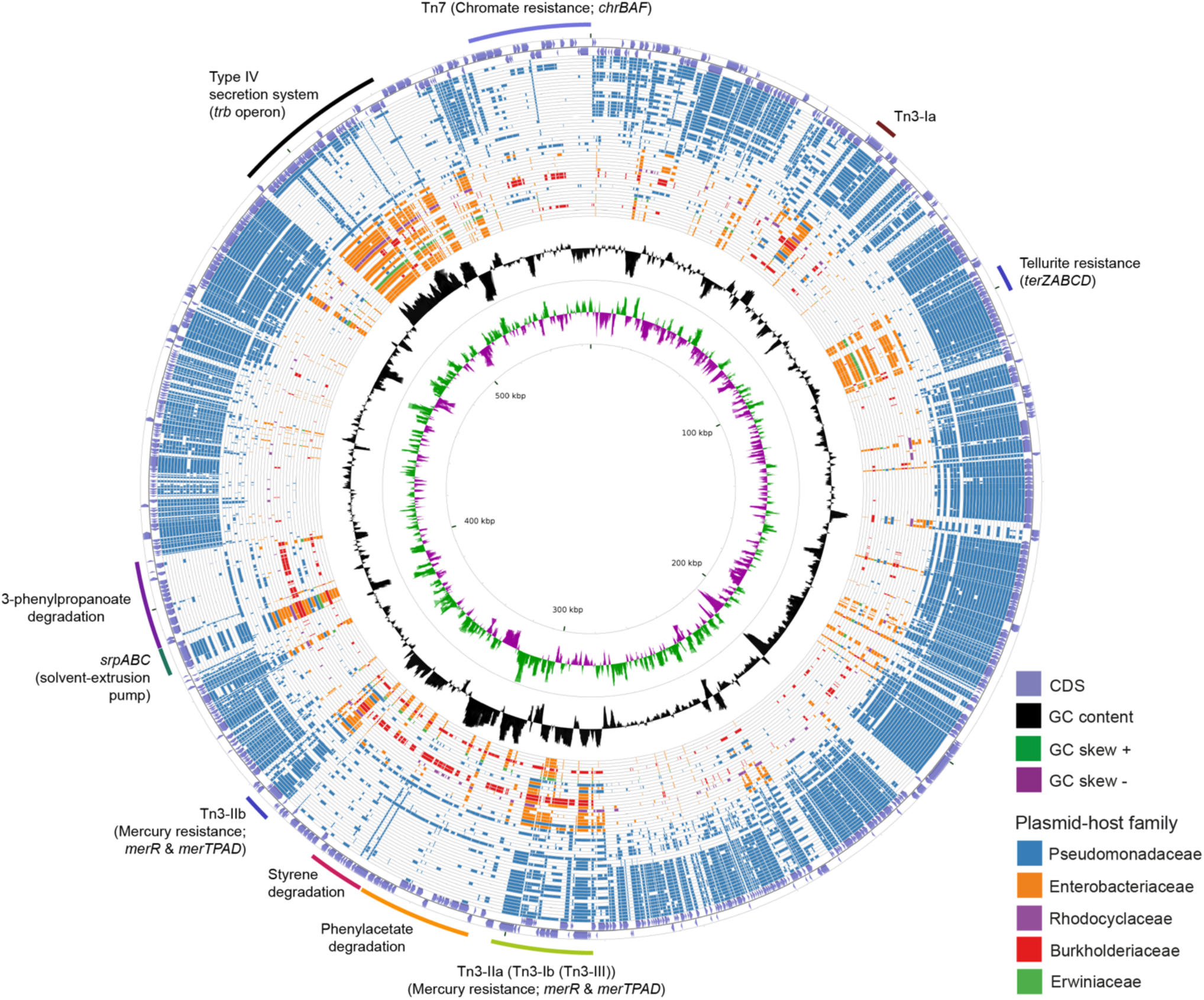
Circular plot of the top 50 plasmids with the highest identity scores to pTTS12. pTTS12 coding sequences (CDS) were aligned to 28915 other plasmids available at NCBI Refseq and Nuccore databases. The outer light purple ring represents the CDS of plus and minus strand of pTTS12. In the inner ring, the GC content is represented in black and the positive and negative GC skew are represented in green and purple respectively. The plasmids are ordered based on their similarity and coverage to pTTS12 CDS from outermost to innermost of the plot with each ring representing a single plasmid as listed in Table 1. Ring colors represent different plasmid-host families; Pseudomonadaceae (blue), Enterobacteriaceae (orange), Rhodocyclaceae (purple), Burkholderiaceae (red), and Erwiniaceae (green). The position of the gene clusters of interest are annotated in the figure. An extended circular plot of the top 500 plasmids with highest identity scores to pTTS12 are shown in Figure S1.

The majority of plasmids highly similar to pTTS12 were found in *Pseudomonas* type strains (Table 1). The relative identity scores presented in Table 1 were calculated as a percentage of total scores of each plasmid divided by the total identity score obtained for pTTS12 itself. The plasmid most similar to pTTS12 is pOZ176 from *P. aeruginosa* PA96 which was previously categorized as a member of incompatibility group P-2 (IncP-2). pOZ176 and pTTS12 share no less than 71% similarity of their encoded genes indicating that these plasmids share a similar IncP-2 backbone. However, while pOZ176 encodes a carbapenem-resistance gene cluster typical for several pathogenic *P. aeruginosa* strains (2), pTTS12 of *P. putida* S12 does not share this feature.

Surprisingly, pTTS12 shares only 21% of sequence similarity with the pSTY plasmid of *P. taiwanensis* VLB120. The major and only gene clusters shared between pTTS12 and pSTY are involved in styrene degradation, phenylpropionic acid degradation, and solvent efflux pump (SrpABC). Interestingly, the genes shared between pTTS12 and pSTY are absent in all other *Pseudomonas* plasmids, except for the solvent efflux pump gene cluster which is similar with TtgGHI efflux pump from the pGRT-1 plasmid of *P. putida* DOT-T1E. The regions encoding solvent efflux pump and phenylpropionic acid degradation are clustered together and have identical synteny in pTTS12 and pSTY (Figure S2). However, the plasmids do not share any mobile genetic elements surrounding this cluster that might indicate a mechanism of acquisition or transfer. The similarity of the styrene degradation clusters and solvent efflux pump - phenylpropionic acid degradation clusters between pTTS12 and pSTY were 99% and 80%, respectively.

Between 8% to 12% of encoded proteins from pTTS12 can also be found in other genera than *Pseudomonas* (Table 1). Several plasmids from *Salmonella, Enterobacter, Citrobacter, Leclersia, Klebsiella, Pantoea, Polaromonas*, and *Cupriavidus* species shared homologous proteins involved in the T4SS conjugation, replication machinery, and plasmid maintenance. Like other IncP-2 plasmids, pTTS12 contains multiple transposable elements, particularly Tn3 mobile elements (Figure 1). This mobile element encodes for heavy metal resistance genes that are predominantly present in IncP-2 plasmids. Additionally, pTTS12 carries a unique Tn7-element consisting of several chromate resistance genes.

### Distinctive conjugation machinery of pTTS12

Megaplasmid pTTS12 contains a P-type T4SS (Type IV secretion system) conjugation system (RPPX_28670-RPPX_28725), sharing synteny with the prototype *trb* operon from *A. tumefaciens* pTiC58 (Figure 2A). For an operational T4SS conjugation machinery, a T4SS gene cluster, type IV coupling protein (T4CP) and a relaxase protein are required (32). An additional *traG* (RPPX_28650), which may serve as a T4CP, is located upstream of the T4SS cluster. Further upstream of the T4SS and T4CP, a putative virD2-like gene (RPPX_28750) and an operon consisting of *parB, parA* and *repA* (RPPX_28765-28775) are encoded on pTTS12. The putative virD2-like gene (RPPX_28750) may play a role as a relaxase which is important for the transferability of pTTS12.

**Figure 2.**
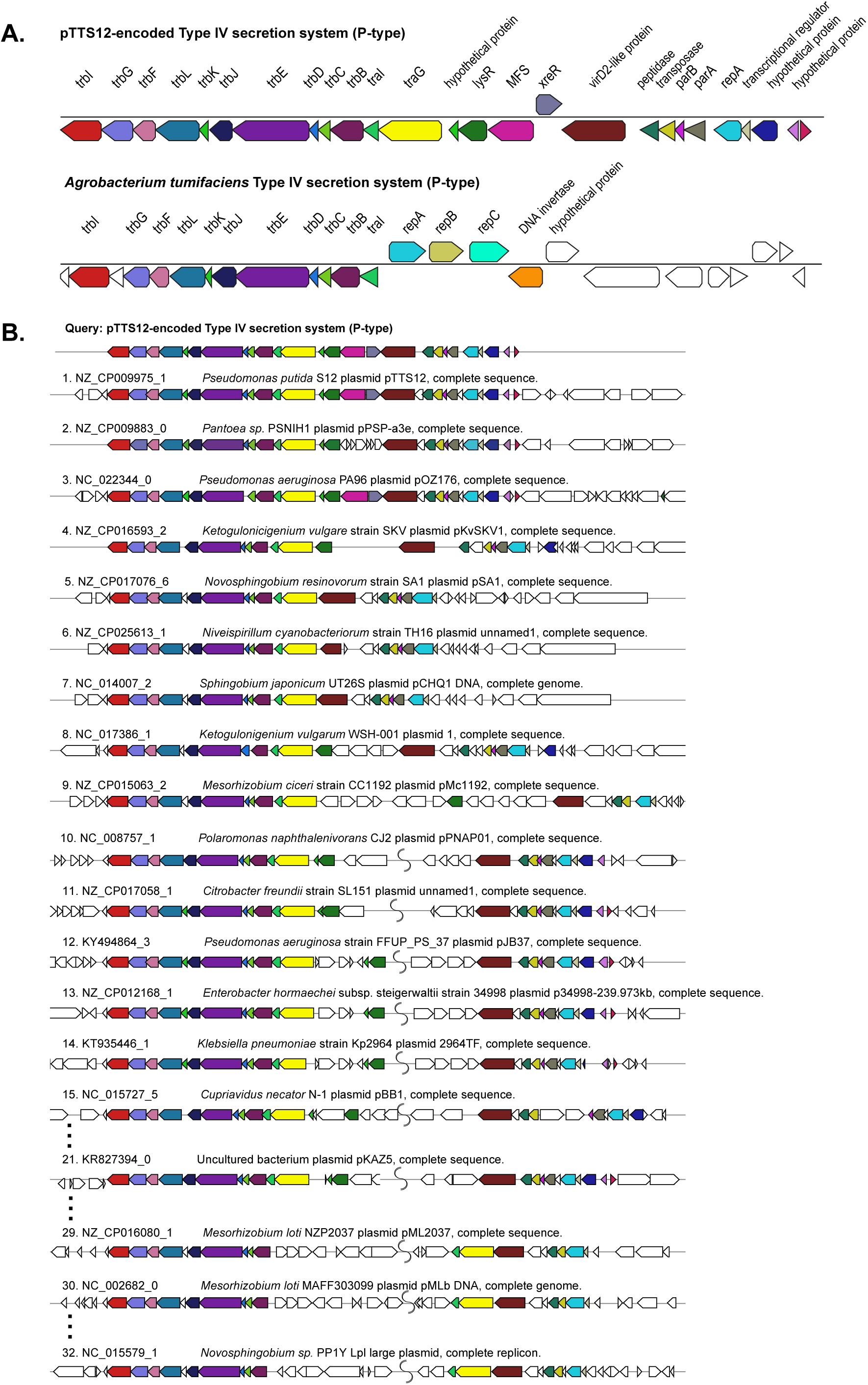
Structure and synteny of pTTS12 conjugation system. A. The arrangement of the T4SS gene cluster found in pTTS12 and the prototype *trb* operon from *Agrobacterium tumefaciens* (pTiC58). The colors represent different genes in the cluster and same colors are assigned for the homologous genes. The gene names are indicated above the respective clusters. pTTS12/T4SS and the *trb* operon of pTiC58 share synteny for 11 genes (*trbI* to *traI*), while other parts are clearly different. B. Synteny plot of the T4SS gene cluster of pTTS12 for plasmid conjugation, replication and partitioning compared with other plasmids. This visualization was generated using multigeneblast software (49). The numbers refer to the order of decreasing synteny. For the sake of clarity, several plots were removed from this figure, indicated by the dots. The colors represent different genes in the cluster corresponding to color-coding in panel A. Putative coupling-protein (T4CP) *traG* is indicated in yellow and the putative relaxase *virD2*-like protein is indicated in brown.

T4SS, *traG*, and upstream genes involved in replication and partitioning shared synteny with the T4SS clusters on other plasmids such as *Pantoea sp*. PSNIH1 (pPSP-a3e), *Pseudomonas aeruginosa* PA96 (pOZ1760), and *Ketogulonicigenium vulgare* SKV (pKvSKV1) (Figure 2B). Typically, *traG* (T4CP) is coupled to the same operon of T4SS while the region encoding replication and partitioning may be separated (Figure 2B), in some cases relatively far. It is interesting to note that in two *Mesorhizobium loti* strains and *Novosphingobium sp*. PP1Y, the T4SS operon and *traG-traI* are completely separated (Figure 2B). Instead, VirD2-like protein (relaxase), *traG* (T4CP) and *traI* formed a single operon.

Most of the Pseudomonadaceae-family plasmids do not contain this *trb* operon (Figure 1) except for pTTS12, pOZ176 from *P. aeruginosa* PA96, pJB37 from *P. aeruginosa* FFUP_PS_37 and an unnamed plasmid from *P. aeruginosa* PA83. Therefore, this conjugative operon may not be common in IncP-2 plasmid family. However, this *trb* operon is abundant among Enterobacteriaceae-family plasmids (Figure 1).

### Distribution of transposable elements conferring heavy metal resistance

pTTS12 as well as many other highly similar plasmids contain a multitude of mobile genetic elements, such as ISS12 and Tn3-family transposases. pTTS12 harbors five copies of Tn3-family transposable elements; two copies of Tn3-I with highest identity to Tn4656, two copies of Tn3-II that are highly similar to Tn5053 and a single copy of Tn3-III with highest similarity to Tn5563 (Figure 3A). Tn3-I is an identical transposase (RPPX_RS27515, *tnpR* and RPPX_RS27515, *tnpA*) to Tn4656 encoded on pWW53 plasmid from *P. putida* MT53. Moreover, the 39-bps inverted repeat (IR) sequences found on both ends of these two elements are highly identical with only a single mismatch difference. In addition to *tnpA* and *tnpR*, this Tn3-element contains a putative methyl-accepting chemotaxis protein (*mcpT*-2) and an insertion sequence IS256. Tn3-II of pTTS12 is identical (99.8% similarity) to Tn5053 of *Xanthomonas sp*. W17 encoding a mercury resistance gene cluster *merR* and *merTPAD* (7). This Tn3-II has 25 bp inverted repeats, bracketing the ends and 5 bp directed repeats (DR). Tn3-III is identical to Tn5563, that is found in several other plasmids, eg. pAMBL of *P. aeruginosa*, pSTY of *P. taiwanensis* VLB-120 and pRA2 of *P. alcaligenes*. The Tn3-III has identical 38 bps IRs bracketing the element, also identical to IRs found for this element in other plasmids. This element contains several genes such as *merR, merTP, pilT* and a gene with a PIN nuclease domain.

**Figure 3.**
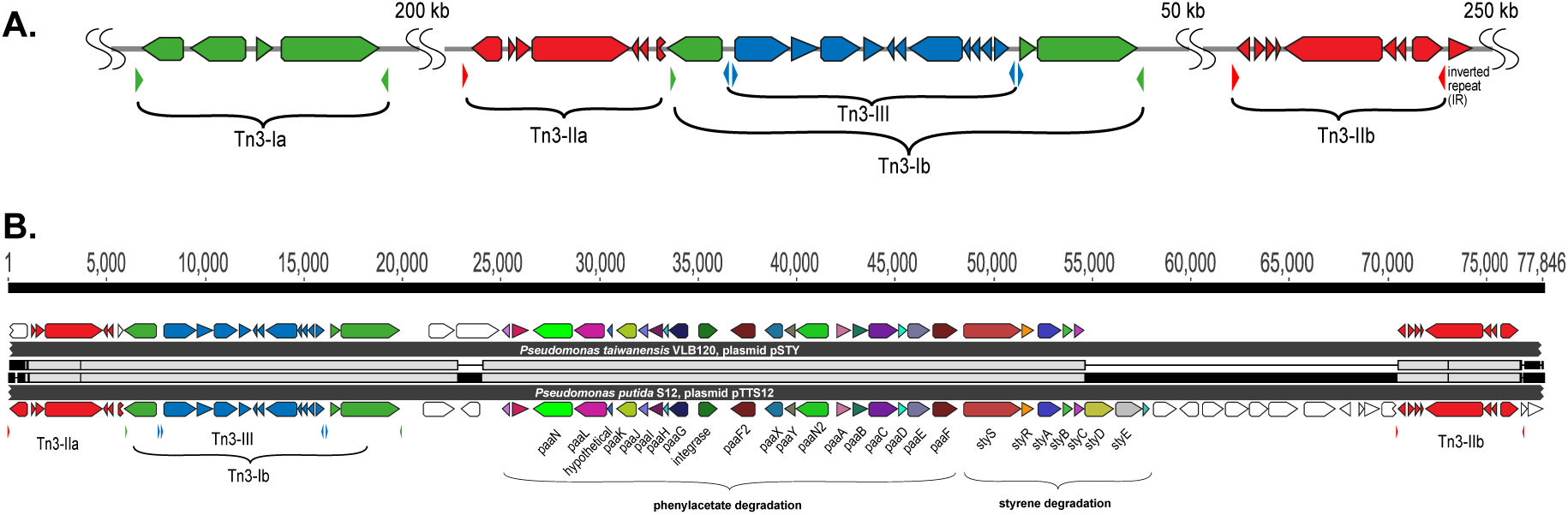
Tn3 transposon family elements responsible for horizontal gene transfer of styrene and phenylacetate degradation pathway. A. Genetic organization of the three different Tn3-family transposable elements in pTTS12. Colors represent the three different Tn3 transposon families; Tn3-I with highest identity to Tn4656 (green), Tn3-IIa and b with highest identity to Tn5053 (red) and Tn3-III with highest identity to Tn5563 (blue). The inverted repeats (IRs) flanking each element are marked by small lowered triangles in the same colours, accordingly. B. Alignment of the Tn3-mediated horizontal gene transfer of the styrene and phenylacetate degradation cluster in pTTS12 (bottom) and pSTY from *Pseudomonas taiwanensis* VLB120 (top). Colors represent the different genes constituting styrene-phenylacetate degradation cluster and the three Tn3 transposable elements, corresponding to the color-coding in panel A. Gene names are indicated.

The Tn4656 and Tn5563 are both duplicated and rearranged on the megaplasmid together with Tn5053. This rearrangement resulted in a highly characteristic sequence, which partially resulted from insertion of Tn5053 transposition in the second copy of Tn4656 (Tn4656-II), between the *mcpT*-2 and *tnpR* loci (Figure 3). The IRs as well as DRs of Tn5053 indicate the insertion site of this element. The IRs of Tn4656-II are well preserved bracketing both elements. Other parts of this sequence consist of the second copy of Tn5563 (Tn5563-II), which is truncated at the right end by insertion of Tn4656-II in *merT* and thus truncated the 5’ end of *merT* sequence, can be identified. Due to this truncation, the IRs of Tn4656-II cannot be found on the right side of the element anymore except for the right IR of initial Tn4656, which is located 58 kbps upstream of this element.

The unique rearrangement of the three different Tn3 elements in pTTS12 surprisingly is also present on pSTY of *P. taiwanesis* VLB120, including an additional copy of Tn5563. In between these two Tn5563-elements, both megaplasmids contain the complete styrene degradation pathway, enabling both strains to grow on styrene as sole carbon source (25, 30). Detailed comparison of the regions between the two Tn5563 elements from pTTS12 and pSTY reveals a high similarity between Tn5563-I and Tn5563-II (Figure 3B). This sequence is around 77 kbps and 60 kbps for pTTS12 and pSTY respectively, including the IRs bracketing both sequences.

In addition to Tn3-elements, a unique Tn7-like element (20 kbps) is also encoded on pTTS12 (Figure 1). Interestingly, an identical transposable element is also encoded on the chromosome of *P. putida* S12. This element contains several transposase genes at both ends and a putative chromate resistance gene cluster (locus tag RPPX_28930-29030; see Table S1). The putative chromate resistance gene cluster was identified based on homology search using BLAST and alignments to other plasmids. RPPX_28995, RPPX_28990, and RPPX_29000 were identified to encode for a chromate resistance efflux pump ChrA, and two regulatory proteins ChrB and ChrF respectively similar to the chromate resistance gene cluster carried by pMOL28 of *Cupriavidus metallidurans* CH34 (33).

### Conjugative megaplasmid pTTS12 is highly stable in *P. putida* KT2440

To characterize the conjugative transferability of pTTS12, we performed biparental mating between *P. putida* S12.1 and *P. putida* KT-BG35. *P. putida* KT-BG35 is a strain derived from *P. putida* KT2440, does not harbour a megaplasmid and carries a gentamicin resistant marker and green fluorescence protein (GFP) at its *attn*7 site (34). The transfer was performed using biparental mating between *P. putida* S12.1 containing pTTS12 with kanamycin resistant marker and *P. putida* KT-BG35. Transconjugant colonies resistant to both kanamycin and gentamicin occurred at the frequency of 4.20 (± 0.51) x 10^−7^. After appropriate selection on agar plates, the identity of transconjugant colonies was confirmed by observing GFP expression of the colonies derived from *P. putida* KT-BG35. Additionally, transfer of entire pTTS12 was confirmed using PCR. Amplification of several regions of pTTS12 using primer pairs 53,496_Fw-56,596_Rv, 200,497_Fw-203,602_Rv, and 286,448_Fw-289,462_Rv resulted in an expected band of 3 kbp in transconjugant colonies. All 15 randomly selected colonies chosen for colony PCR showed correct bands. This confirmed the transfer of entire pTTS12 into *P. putida* KT-BG35 and the resulting strains will further be referred to as *P. putida* KTpS12. Several attempts to transfer pTTS12 into *E. coli* strains represented by *E. coli* MV1190 and *E. coli* XL1-Blue by biparental mating did not result in any successful transconjugant colonies.

The putative relaxase gene, *virD2*, was identified on pTTS12 (locus tag RPPX_28750). To confirm this finding, we created a complete deletion of *virD2* gene and compared the self-transfer frequency of pTTS12 Δ*virD2* and wild-type pTTS12 into *P. putida* KT2440. The transfer frequency of pTTS12 Δ*virD2* was 4.16 (± 0.08)× 10^−7^, which was not significantly different (p-value 0.8867) compared to the transferability of the wild-type pTTS12 from *P. putida* S12 to *P. putida* KT2440.

The stability of pTTS12 in *P. putida* KTpS12 and *P. putida* S12 was examined for 5 passages in the absence of antibiotic selective pressure (approximately 10 generations/passage step). No plasmid loss was observed from either *P. putida* KTpS12 or *P. putida* S12 growing without selective pressure while the negative control pSW-2 showed a steady plasmid loss (Figure 4A). Serendipitous deletion of pTTS12 was not found in *P. putida* S12 and *P. putida* KTpS12 either. Hence, pTTS12 is a stable megaplasmid in both *P. putida* S12 and *P. putida* KT2440.

**Figure 4.**
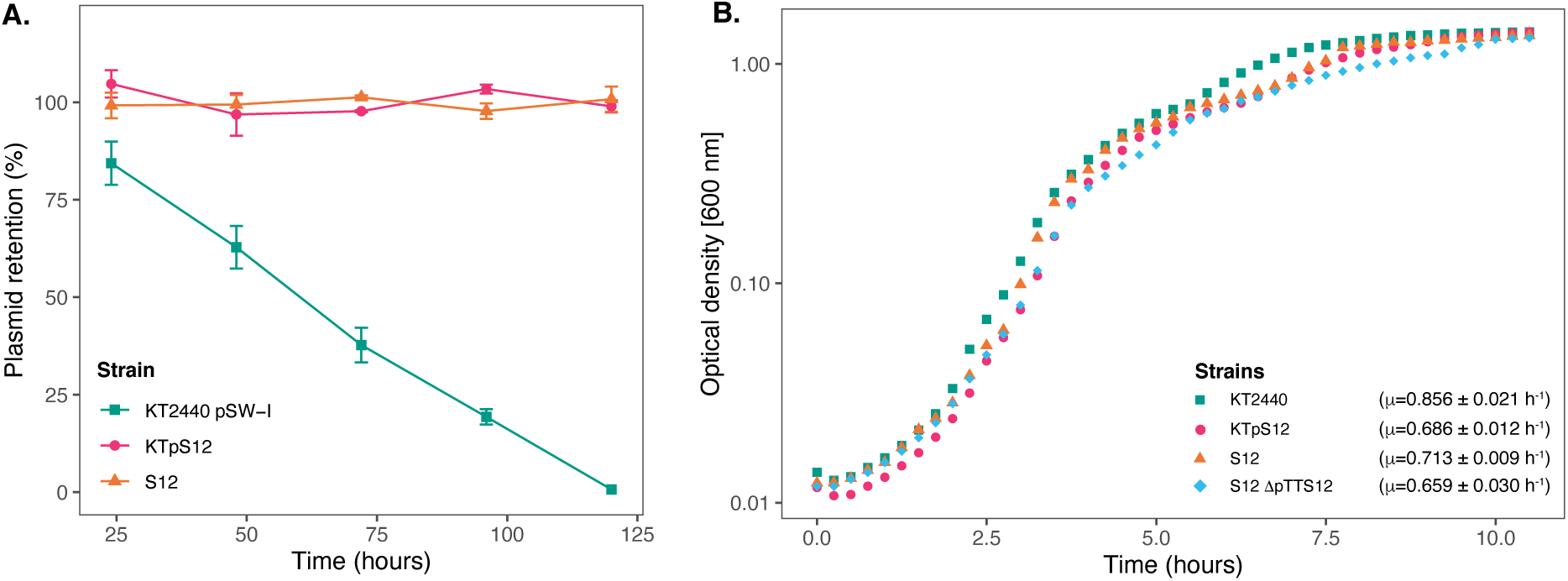
Megaplasmid pTTS12 is highly stable in *P. putida* strains and reduces maximal growth rate in *P. putida* KT2440. A. Plasmid retention of pTTS12 in *P. putida* KT2440 and *P. putida* S12 on liquid LB without selection pressure for approximately 50 generations. Colors and shapes indicate the different strains with green squares indicating *P. putida* KT2440 pSW-II, magenta circles indicating *P. putida* KTpS12, and orange triangles indicating *P. putida* S12 (containing pTTS12). Plasmid pSW-II in *P. putida* KT2440 was used as a control for loss of unstable plasmid. pTTS12 is stably maintained in both *P. putida* S12 and *P. putida* KTpS12. B. Growth curve of *P. putida* KT2440, *P. putida* KTpTTS12 *P. putida* S12, and *P. putida* S12 ΔpTTS12. Growth was followed on liquid LB at 30 °C, 200 rpm shaking. Growth curves represent data obtained from three biological replicates for each strain, starting from OD_600nm_ = 0.05. Green squares indicate *P. putida* KT2440 pSW-II, magenta circles indicate *P. putida* KTpS12, orange triangles indicate *P. putida* S12 (containing pTTS12), and blue diamonds indicate *P. putida* S12 ΔpTTS12. Maximum growth rates calculated for each strais are indicated within the figure.

The occurrence of large plasmids in bacteria is known to cause metabolic burden, which is reflected in reduced bacterial growth rate (μ). *P. putida* KTpS12 exhibited a lower maximum growth rate compared to *P. putida* KT2440, 0.686 ± 0.012 and 0.856 ± 0.021 h^-1^ respectively (Figure 4B). However, *P. putida* S12 and *P. putida* S12 ΔpTTS12 did not show a significant maximum growth rate difference (0.713 ± 0.009 and 0.659 ± 0.030 h^-1^ respectively). The length of lag-phase and biomass yield at stationary phase remained unaffected with pTTS12 present in both *P. putida* S12 and KTpS12. Apparently, the presence pTTS12 imposes a only slight metabolic burden in *P. putida* KT2440, but not in S12.

### Phenotypic features of pTTS12 and attained solvent tolerance in *P. putida* KT2440

*P. putida* S12 is intrinsically tolerant to an assortment of environmental xenobiotics. The majority of these characteristic traits are encoded on pTTS12; the styrene degradation operon *styABCDE*, the tellurite resistance operon *terZABCDE* and a solvent efflux pump *srpABC*. To explore functionality of pTTS12 following conjugative transfer, these characteristic traits were investigated in *P. putida* KTpS12 (Figure 5). Activity of the styrene degradation operon in *P. putida* KTpS12 was tested by growing on minimal media with styrene as carbon source and inspecting transformation of indole into indigo. Similarly as observed in *P. putida* S12, *P. putida* KTpS12 was able to transform indole into indigo, whereas wild-type KT2440 was not, indicating activity of styrene monooxygenase (*styA*) and styrene oxide isomerase (*styB*) in these strains (Figure 5A). In addition, *P. putida* KTpS12 was able to grow on minimal media supplemented with styrene as carbon source or minimal media agar plates incubated in styrene atmosphere (data not shown).

**Figure 5.**
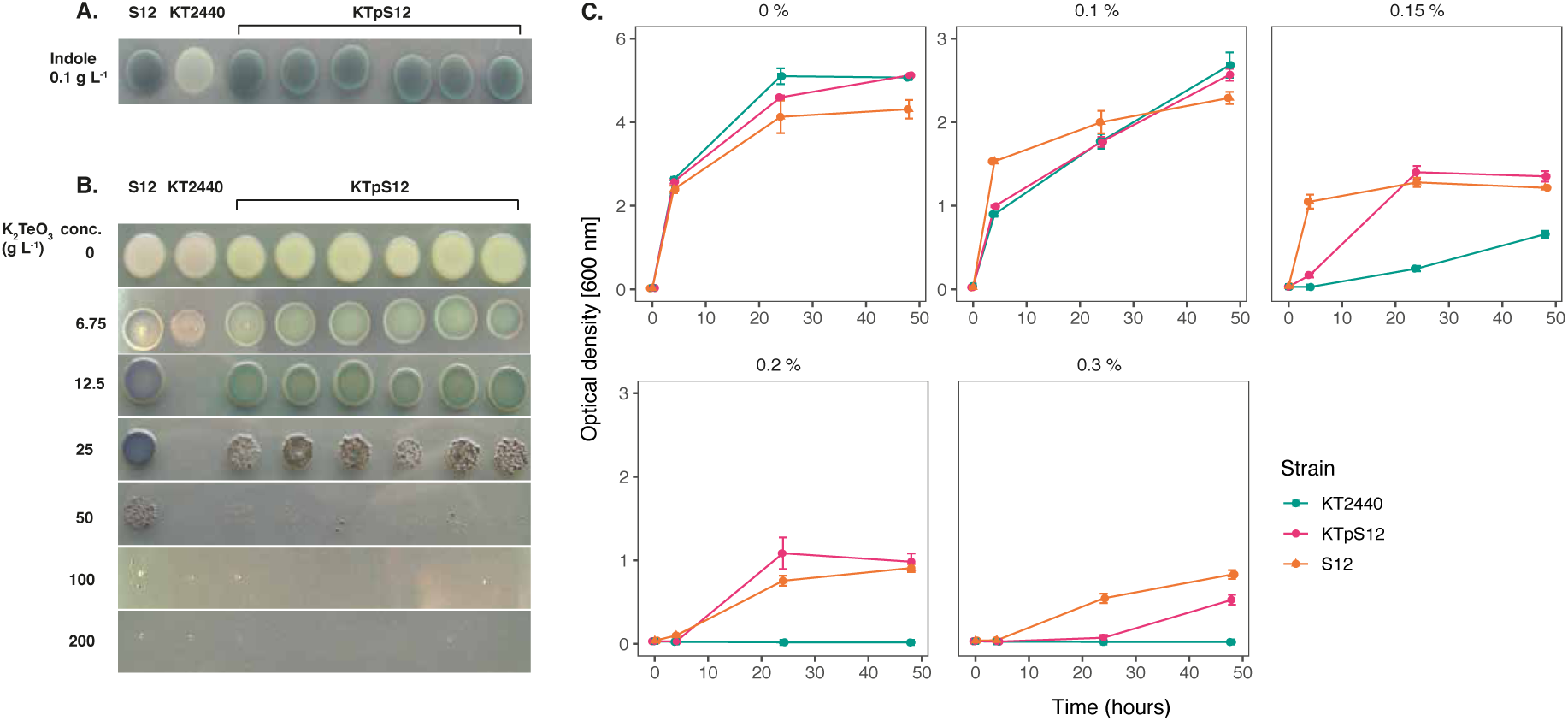
Transfer of pTTS12 phenotypic characteristics. A. Production of indigo from indole indicating activity of styrene monooxygenase (StyA) and styrene oxide isomerase (StyB). Minimal medium (M9) was supplemented with indole which, in the presence of StyAB enzymes encoded on pTTS12, is converted into indigo. Positive control *P. putida* S12 showed indigo coloration, whereas negative control *P. putida* KT2440 remained white. *P. putida* KTpS12 showed indigo coloration indicating activity of pTTS12-encoded *styAB*. B. Growth of *P. putida* strains in the presence of potassium tellurite (K_2_TeO_3_). Minimum inhibitory concentration (MIC) of positive control *P. putida* S12 was 200 g L^-1^ whereas the MIC of the wild-type *P. putida* KT2440 was 12.5 g L^-1^. The MIC of *P. putida* KTpS12 was 100 g L^-1^ indicating the presence and activity of the *ter* operon on pTTS12. C. Growth of *P. putida* strains on increasing concentrations of toluene. Optical density (OD_600nm_) of the cultures was measured at 4, 24, and 48-hour time points with starting point of OD_600nm_ = 0.1. The y-axis range is different between the first panel (0-6) and the other panels (0-3). Green squares indicate *P. putida* KT2440 pSW-II, magenta circles indicate *P. putida* KTpS12, and orange trianglse indicate *P. putida* S12 (containing pTTS12). *P. putida* KTpS12 showed an increase in solvent tolerance indicating the presence and acivity of the *srp* operon from pTTS12.

Potassium tellurite (K_2_TeO_3_) exhibits antimicrobial activity and tellurite resistance in bacteria is achieved through reduction of tellurite (TeO_3_^2-^) into a less toxic metallic tellurium (Te^0^), hence the formation of black colonies in the presence of tellurite (35). The resistance genes on pTTS12 are encoded as an operon *terZABCDE* (RPPX_26360-26385). To investigate whether this gene cluster is able to increase resistance towards tellurite in *P. putida* KT2440, the minimum inhibitory concentration (MIC) was determined for the three different stains (Figure 5B). The MIC of potassium tellurite for *P. putida* KT2440 was 16 fold less compared to *P. putida* S12, 12.5 mg L^-1^ and 200 mg L^-1^ respectively. Interestingly, *P. putida* KTpS12 showed an 8 fold increase in tellurite resistance (MIC 100 mg L^-1^) compared to its parental strain *P. putida* KT2440.

The solvent efflux pump *srpABC* encoded on the megaplasmid pTTS12, enables survival and growth of *P. putida* S12 in non-utilized organic solvents (36). In order to investigate the effect of induced tolerance toward organic solvents due to the introduction of pTTS12 in *P. putida* KTpS12, a growth assay was performed in the presence of different toluene concentrations (Figure 5C). *P. putida* KT2440 was able to grow in LB liquid media supplemented with toluene up to a maximum of 0.15 % v/v concentration, although suffering from a significant growth reduction compared to *P. putida* S12. With the introduction of pTTS12, *P. putida* KTpS12 tolerance toward toluene increased to 0.30 % v/v. Similar concentration was obtained for *P. putida* S12, while *P. putida* KTpS12 exhibited slightly slower growth in the presence of toluene. These observations indicated that the introduction of megaplasmid pTTS12 provided a full set of characteristic features to *P. putida* KT2440.

## Discussion

### Megaplasmid pTTS12 defines an environment-adaptive and type-specific family of IncP-2 plasmids, carrying distinct accessory gene clusters

Megaplasmid pTTS12 of *P. putida* S12 is closely related to other plasmids in proteobacteria, especially within the *Pseudomonas* genus. In this study we demonstrated high similarity of the pTTS12 ‘backbone’ with pOZ176 from *P. aeruginosa* PA96 (2) and several other IncP-2 plasmids, such as pQBR103 of *Pseudomonas fluorescens* SBW25 (37). pOZ176 has been categorized within incompatibility group IncP-2 based on its replication, partitioning, and transfer machinery along with conferring tellurite resistance as a key feature of IncP-2 plasmid (2, 5). Indeed, pTTS12 encodes typical characteristics of IncP-2 plasmids, such as heavy metal resistance (tellurite, mercury, and chromate) and plasmid maintenance via the *parA*/*parB*/*repA* system (10, 38). pTTS12 clearly conferred tellurite resistance when transferred to tellurite-susceptible *P. putida* KT2440, confirming this IncP-2 characteristic phenotype. Interestingly, despite high similarity between pTTS12 and pOZ176, pTTS12 lacks the Tn6016 *bla*_IMP-9_-carrying class 1 integron cassette which is an important trait of pOZ176 conferring resistance to aminoglycosides and carbapenems (2, 5). On the other hand, pOZ176 lacks the solvent efflux pump, styrene degradation pathway, and phenylpropionic acid degradation pathway which are main characteristics of pTTS12. This demonstrates that very similar megaplasmid backbones may carry resistance to a diversity of xenobiotics, metabolic functions, or virulence gene clusters (39). Whereas the traits disseminated by pTTS12 and pOZ176 respectively, are highly divergent and distinctive, a similar observation of environment-adaptive traits conferred by flexible and diverse accessory gene clusters contained on IncP-2 plasmids was recently reported for *P. aeruginosa* plasmids pBT2436 and pBT2101, carrying multiple MDR cassettes (1).

### Convergent distribution of plasmid-encoded solvent tolerance gene clusters

pTTS12 shares its unique features of styrene and phenyl-propanoate degradation pathway with *P. taiwanensis* VLB120 and the solvent extrusion pump SrpABC with TtgGHI from *P. taiwanensis* VLB120 and *P. putida* DOT-T1E. These solvent tolerant strains were isolated independent of each other, although in different geographical locations; *P. putida* S12 and *P. taiwanensis* VLB120 were isolated for their ability to utilize styrene as sole carbon source (11, 40). *P. putida* DOT-T1E was isolated for its ability to degrade toluene (9). Hence, these plasmids isolated from different environmental sources show convergent distribution of highly similar gene clusters with similar features related to environmental stresses (e.g. organic solvents) on different plasmid backbones. In *P. taiwanensis* VLB120, the arrangement of gene clusters encoding the styrene degradation pathway and solvent efflux pump - phenylpropionic acid degradation pathway is highly similar to pTTS12, with 99% and 80% similarity, respectively. This moreover, indicates that exchange of the styrene degradation pathway occurred more recently than exchange of the efflux pump/phenylpropionic acid degradation cluster. Alternatively, these clusters may have been acquired independently.

The styrene degradation gene cluster is encoded within a unique arrangement of Tn3 family transposases shared by pTTS12 and pSTY from *P. taiwanensis* VLB120 (Figure 3). Due to the transposition of Tn3-Ib, Tn3-IIa lost its right flank inverted repeat (IR). Because of this arrangement, Tn3-IIa can only “jump” while carrying the entire Tn3 and styrene-phenylacetate degradation clusters using the right flank of Tn3-IIb IR. This may explain the occurrence of the styrene-phenylacetate degradation cluster distinctive arrangement shared between pTTS12 and pSTY. Although we could not find evidence of mobile genetic elements carrying the efflux pump and the phenylpropionic acid degradation gene clusters on pSTY and pTTS12, it is well possible that the exchange originally occurred via such route.

### Transferability of pTTS12

pTTS12 contains a P-Type type IV secretion system (T4SS) with synteny similar to the prototype *trb* operon of the *A. tumefaciens* pTiC58 system (32, 41). Indeed, our experiments showed that pTTS12 is transferable towards other *P. putida* strains with a frequency of 4.20 (± 0.51) × 10^−7^. A putative relaxase, responsible for creating a nick prior to plasmid transfer via the conjugative bridge, was predicted to be encoded by *virD2* (locus tag RPPX_28750). However, complete deletion of *virD2* did not resulted in a significant reduction of transfer frequency, compared to the wild-type pTTS12. This may indicate the presence of other relaxase(s) which act in-trans with the P-type T4SS to mediate the nicking of plasmid DNA and subsequently aided the plasmid transfer process. *P. putida* S12 contains other T4SS within its genome which share synteny with the I-type T4SS represented by Dot/Icm from *L. pneumophila* (Figure S3), although it is unclear whether Dot/Icm can support the transfer of pTTS12 (42).

### Amelioration of pTTS12 reduces fitness cost and promotes plasmid persistence in *P. putida* S12

We observed that in its original host *P. putida* S12 on rich media, pTTS12 did not impose an apparent metabolic burden whereas in another strain as *P. putida* KT2440, pTTS12 caused a 20% reduction of maximum growth rate. We previously reported on the occurrence of metabolic burden caused by pTTS12 in *P. putida* S12 in the presence of toluene (43). In addition to the relatively low metabolic burden, pTTS12 was very stable in both *P. putida* S12 and *P. putida* KT2440 (Figure 4A). Several mechanisms may be involved in the reduced fitness cost and the stability of pTTS12 in *P. putida* S12.

Conjugative transfer may impose substantial cost and burden due to the energy investment on pili formation during conjugation (39, 44). Indeed, pTTS12 showed a substantially lower conjugation rate (4.20 × 10^−7^ transfer frequency) after 24 hours in comparison to other IncP-2 plasmids (10^−1^ to 10^−4^ transfer frequency after 2 hours) (38). Downregulation of plasmid genes appeared to be involved in the amelioration of pTTS12. The expression of plasmid-encoded resistance genes, like the solvent extrusion pump, can typically be a source of metabolic burden imposed by plasmids (45). pTTS12 contains multiple types of mobile genetic elements, with ISS12 being the most abundant (4, 27). Substantial duplication of the ISS12 mobile element has previously been reported to interrupt *srpA*, encoding the periplasmic subunit of the solvent efflux pump (4). In the prolonged absence of organic solvent, expression and maintenance of the solvent efflux pump may be costly for the bacterial cell, hence, interruption of the *srp* efflux pump gene cluster may further reduce the plasmid burden of pTTS12. In addition to these mechanisms, we recently described the contribution of a toxin-antitoxin module SlvT-SlvA to the stability of pTTS12 (43).

### Future outlook

IncP-2 family plasmids are widely distributed among environmental and clinical *Pseudomonas* isolates. These plasmids contain variable regions encoding MDR, xenobiotics extrusion pumps, degradation pathways, and heavy metal resistance cassettes. In contrast to the dynamic variable regions, the core backbone of these plasmids shows a general conservation. It is interesting to discover the minimal backbone of IncP-2, which enables Pseudomonads to scavenge gene clusters important for its survival both in environmental and clinical set-up, as a model of horizontal gene transfer. Ultimately, IncP-2 plasmid backbone may be promising for biotechnological and bioremediation applications due to its stability and relatively low metabolic burden. Moreover, these plasmids have been described to exchange their traits, thus creating hybrid plasmids when they occur within the same host (38). The mechanisms for such exchange are poorly understood and may well involve transposition and conjugation as suggested by the shared and type-specific characteristics of pTTS12 and pOZ176 as described in this study. Further clarification of these mechanisms is important to shed light on the rapid dissemination of bacterial tolerance and resistance to antibiotics and chemical stresses in the environment.

Successful attempts have been made in exploiting traits from environmental plasmids for standardized components in synthetic biology (46). Environmental plasmids are exchangeable between different hosts and may express their genetic features in various genomic and metabolic backgrounds. Moreover, they are an excellent source of novel biological parts such as origin of replication, metabolic pathway, resistance marker, and regulated promoters. New sequencing technologies and comparative genomics analyses support the identification of the genes enabling these features. Here, we demonstrated that pTTS12 contains promising exchangeable gene clusters and building blocks to construct robust microbial hosts for high-value biotechnology applications.

## Materials and methods

### Cultivation of P. putida

Strains and plasmids used in this report were listed in Table 2. All *P. putida* strains were grown in Lysogeny Broth (LB) containing 10 g L^-1^ tryptone, 5 g L^-1^ yeast extract and 5 g L^-1^ sodium chloride at 30 °C with 200 rpm shaking. *E. coli* strains were cultivated in LB at 37 °C with 250 rpm shaking in a horizontal shaker (Innova 4330, New Brunswick Scientific). For solid cultivation, 1.5 % (w/v) agar was added to LB. M9 minimal medium used in this report was supplemented with 2 mg L^-1^ MgSO_4_ and 0.2% w/v of citrate as sole carbon source (11). Bacterial growth was observed by optical density measurement at 600 nm (OD_600nm_) using a spectrophotometer (Ultrospec 2100 pro, Amersham Biosciences). Maximum growth rate and other parameters were calculated using growthcurver R-package ver.0.3.0 (47). Solvent tolerance analysis was performed by growing *P. putida* strains in LB starting from OD_600nm_ = 0.1 in Boston bottles with Mininert bottle caps. When required, potassium tellurite (6.75 - 200 mg L^-1^), indole (100 mg L^-1^), gentamicin (25 mg L^-1^), ampicillin (100 mg L^-1^), tetracycline (25 mg L^-1^), and kanamycin (50 mg L^-1^) were added to the media.

**Table 2.**
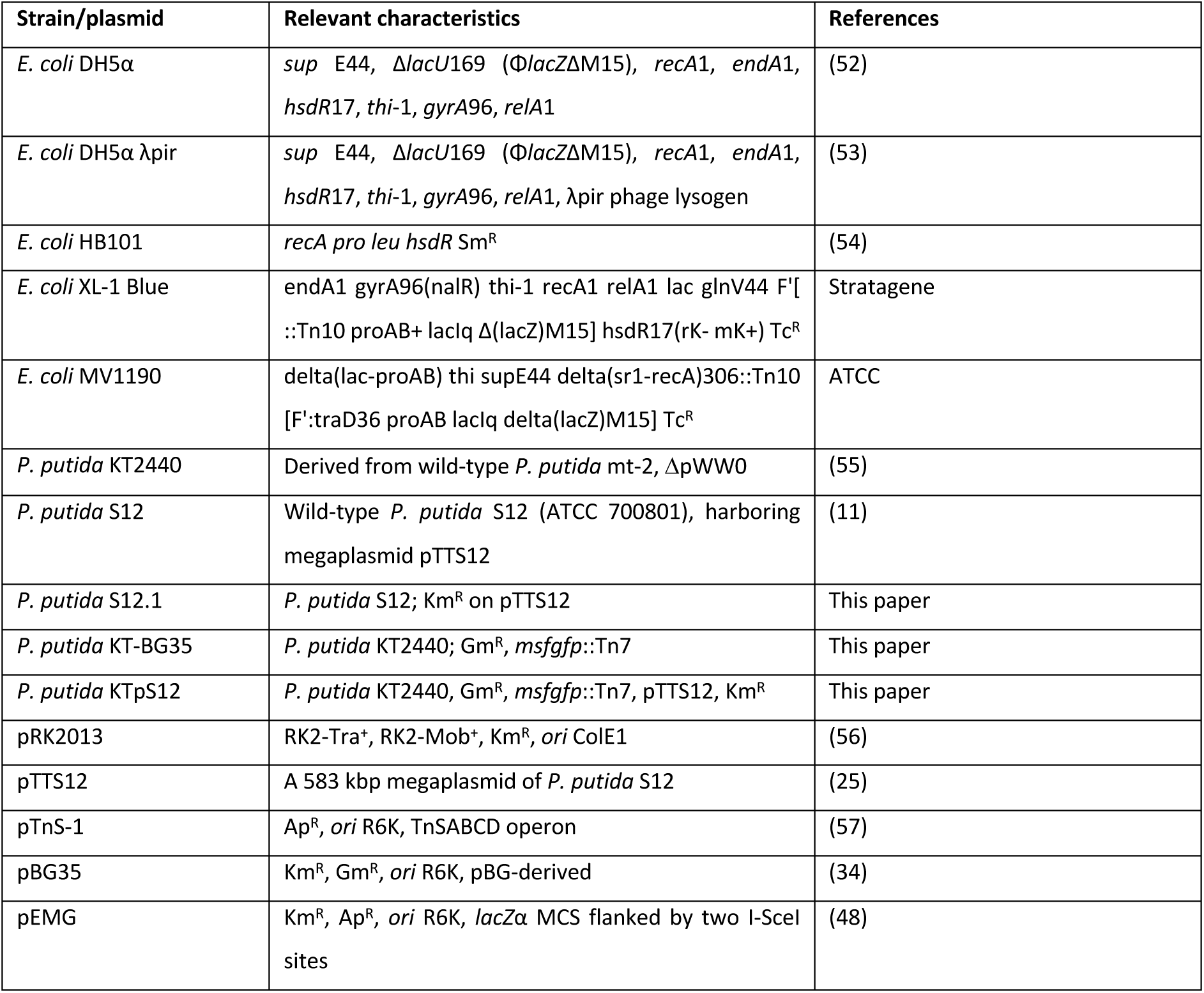
Bacterial strains and plasmids used in this study.

### DNA methods

All PCR reaction were performed using Phire polymerase (Thermo Fischer) according to the manufacturer’s manual. All primers are listed in Table 3 and were obtained from Sigma-Aldrich. PCR reactions were visualized and analyzed by gel electrophoresis on 1 % (w/v) TBE agarose gels containing 5 mg L^-1^ ethidium bromide in an electric field (110V, 0.5x TBE running buffer).

**Table 3.**
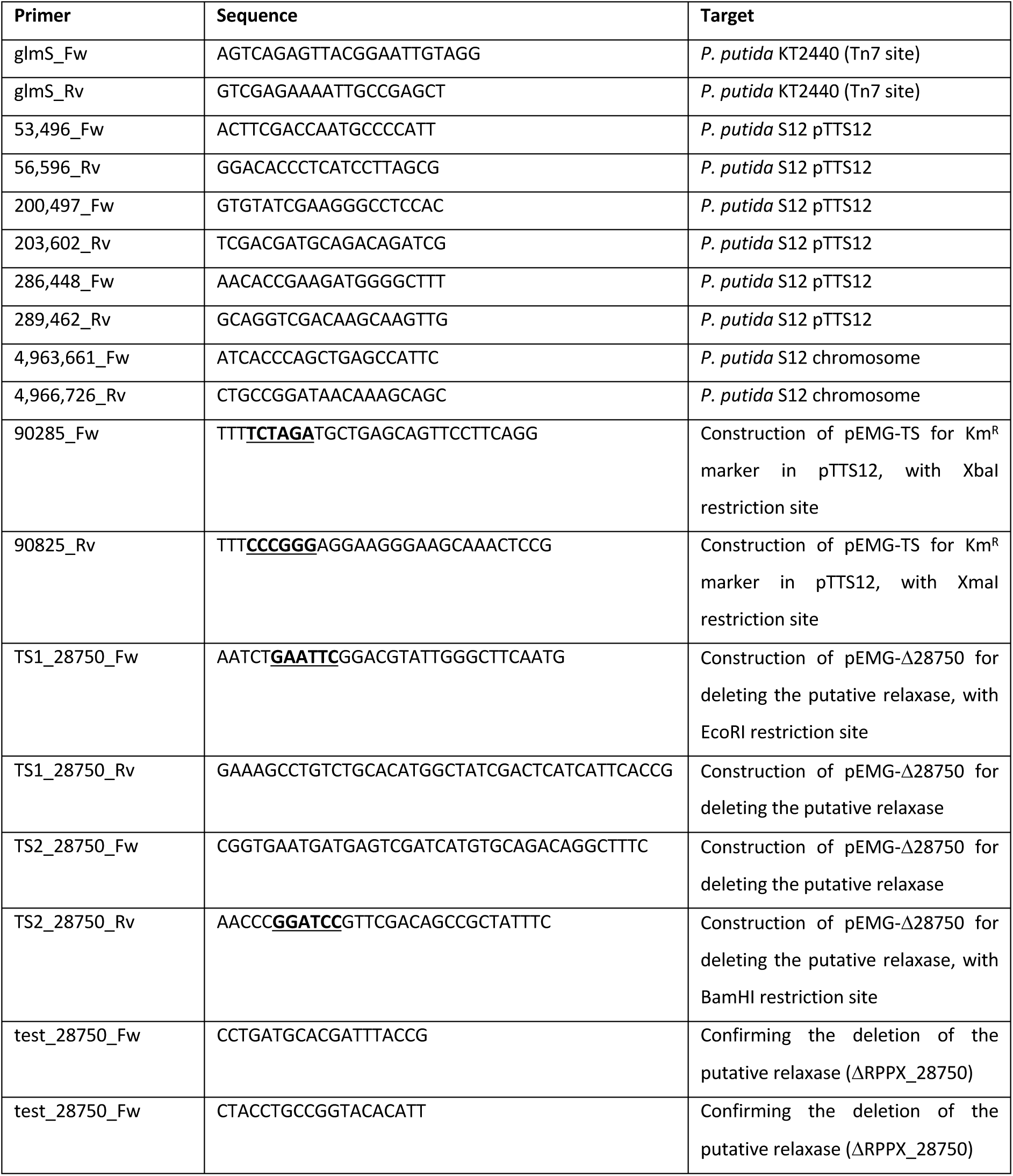
Primers used in this study.

### Megaplasmid pTTS12 transfer into *P. putida* KT2440 and *E. coli* strains

Gentamicin resistance and GFP containing cassette were incorporated into *P. putida* KT2440 chromosome at the Tn7 site using pBG35 plasmid resulting in the strain *P. putida* KT-BG35 as previously described (34). Correct transformants were additionally verified by observing the gentamicin resistance, GFP expression and colony PCR of *P. putida* KT2440 chromosome sequence. Kanamycin resistance gene was introduced into the megaplasmid pTTS12 by integrating plasmid pEMG using homologous recombination resulting in *P. putida* S12.1 as previously described (48). A single homologous recombination site was obtained by PCR with 90285_Fw and 90825_Rv primer pair and this fragment was used to construct pEMG-TS plasmid. Correct integration into *P. putida* S12 was verified by observing kanamycin resistance and colony PCR using primers 4,963,661_Fw-4,966,726_Rv.

The transfer of pTTS12 into *P. putida* KT-BG35, *E. coli* XL1-Blue or *E. coli* MV1190 were performed by biparental mating between *P. putida* S12.1 and the recipient strain on LB agar for 24 hours at 30 °C. The correct transformants were selected using LB agar supplemented with gentamicin and kanamycin for *P. putida* KT-BG35 or with tetracycline and kanamycin for *E. coli* strains. Additionally, transformants were verified using colony PCR (53,496_Fw-56,596_Rv, 200,497_Fw-203,602_Rv, and 286,448_Fw-289,462_Rv). Plasmid transfer rate was determined by comparing the event of successful plasmid transconjugant with the colony formation unit (cfu) of the recipient strain (*P. putida* KT-BG35, *E. coli* XL1-Blue or *E. coli* MV1190) after biparental mating (Gm^R^/Tc^R^). Plasmid stability was determined by calculating the event of megaplasmid loss in *P. putida* KTpS12 grown in liquid media without supplementation of kanamycin as the selective pressure for pTTS12. Plasmid pSW-2 was used as a control for megaplasmid loss events (48).

### Sequence analyses

All plasmid sequences were downloaded from NCBI database, 22389 plasmid sequences from Refseq database (retrieved 8 March 2020) and complemented with few plasmid sequences from NUCORE that were omitted from Refseq. The CGviewer (31) was used to generate circular plots of entire pTTS12 with standard settings and MultiGeneBlast tool (49) was used to generate synteny plots of specific regions and operons. Further analysis was performed using Geneious software (BioMatters), and selected sequences were aligned using MAFFT (50) for DNA sequences or MUSCLE (51) for protein sequences.

## Acknowledgement

We thank Erik Vijgenboom and Gerben Voshol (Leiden University) for helpful discussions and Omar Qachach for his assistance during the data acquisition process. R. Hosseini was funded by the Dutch National Organization for Scientific Research NWO, through the ERAnet-Industrial Biotechnology programme, project “Pseudomonas 2.0”. H. Kusumawardhani was supported by the Indonesia Endowment Fund for Education (LPDP) as scholarship provider from the Ministry of Finance, Indonesia.

